# The molecular basis of pine wilt disease resistance in *Pinus massoniana*

**DOI:** 10.1101/2024.03.13.584545

**Authors:** Jinjin Wang, Shouping Cai, Liqiong Zeng, Jie Tong, Danling Hang, Xinliang Zhang, Yu Fang, Shunde Su, Jun Su

**Author notes:** These authors have contributed equally to this work and share first authorship (Jinjin Wang, Shouping Cai). Correspondence: Shunde Su; Jun Su.

## Abstract

Pine wilt disease (PWD), caused by the pine wood nematode (PWN) *Bursaphelenchus xylophilus*, results in significant economic and ecological damage to *Pinus* forests and plantations worldwide. *Pinus massoniana* is the primary host of PWD in southern China, but its response to the PWN remains largely unstudied. Previously, we observed PWD in a *P. massoniana* nursery that contained over 71 commonly used cultivars. Through field phenotyping, we identified two groups of PWN resistant cultivars. Resistant cultivars (RC) exhibited very low PWN carrying amounts (PCA) and had relatively low mortality, and tolerant cultivars (TC) had high PCA but low mortality. In this study, we confirmed via PWN inoculation assays that the resistant and tolerant cultivars had lower mortality rates, 10% and 11%, respectively, than other cultivars (which had a mortality rate of 83.3%). The RC had a PCA that was significantly lower than that of other cultivars, while the TC exhibited a higher PCA. To explore the molecular mechanisms underlying the response of *P. massoniana* to PWN, the transcriptome and metabolome of the above cultivars were profiled by high throughput sequencing. As no reference genome is available for *P. massoniana*, we generated a new full-length transcriptome library with iso-seq. Using the transcriptome and metabolome from different *P. massoniana* cultivars inoculated with PWN, we found three major PWD resistance strategies. 1) The common response strategy involved three important molecular pathways. First, synthesis of indole-3-acetic acid (IAA) and abcisic acid (ABA) were suppressed during PWN invasion, thus suppressing the synthesis of polysaccharides (especially myo-inositol and trehalose) and (-)-riboflavin through ABC transporters. Second, by inhibiting aspartic acid, the synthesis of arginine and proline through APS5 (aspartate aminotransferase 5) were suppressed. Third, by reducing cysteine, GERD (germacrene D synthase) was up-regulated, and sesquiterpenoid and triterpenoid metabolites were accumulated to resist PWN invasion. 2) The most effective resistance strategy involved the accumulation of reactive oxygen species (ROS), which enhanced jasmonic acid (JA) accumulation and highly induced the expression of chitinase, thus improving resistance to PWN. 3) The tolerance strategy involved the induction of phosphatidylcholine, which promoted flavonoid and anthocyanin synthesis via LPIN (phosphatidate phosphatase), LOX2S (lipoxygenase), and LOX1_5 (linoleate 9S-lipoxygenase), thus significantly inhibiting the pathogenicity of the PWN but not the PCA. These results illustrate the molecular mechanisms by which *P. massoniana* cultivars resist or tolerate PWN and are of great significance in the prevention and control of PWD.

## 1 Introduction

Pine wilt disease (PWD) is one of the most destructive diseases in *Pinus* forests worldwide and poses a serious threat to the health of forest ecosystems in China (Xie et al., 2017; Li et al., 2021; Zhang et al., 2023). *Pinus massoniana* is severely affected by PWD in southern China. Since the disease was first discovered, over 5,000,000 m³of *Pinus* forests have been infected (Cai et al., 2021; An et al., 2023; Li et al., 2023). Due to its hardiness and low maintenance costs, *P. massoniana* is the main cultivated variety of *Pinus* in impoverished areas of southern China, and it is irreplaceable (Yu et al., 2020; Zhu et al., 2022). However, due to the widespread PWD epidemic (Hopf-Biziks et al., 2017; Pajares et al., 2017), planting *P. massoniana* has become a major challenge, and the selection of resistant cultivars is critical to solving this problem.

Cultivating pine wood nematode (PWN) resistant cultivars is currently the most economical and effective approach to reduce PWD and ensure the health of *Pinus* forests (Liu et al., 2022; Hu et al., 2023). Through genetic engineering, traditional breeding, and molecular marker-assisted selection, cultivars can be screened and planted (Li et al., 2010; Török et al., 2019; Thulasinathan et al., 2023). This limits the spread of PWD and enhances the ecological stability, biodiversity, and sustainable development of pine forest ecosystems. In recent years, a number of PWD resistant cultivars have been developed domestically and internationally, such as GD5, DAC12, and AH1 (Deng et al., 2024; Gao et al., 2024). Foreign cultivars such as Clone number 227, Tanabe 54, and Tosashimizu 63 have also exhibited PWD resistance (Hirao et al., 2019; Guo et al., 2023). However, the above cultivars were naturally selected, and their effectiveness in the field is unstable. Genetic engineering can be used to screen resistant cultivars, improve their stability in field production, and significantly reduce the time required for traditional breeding (Li et al., 2010; Mackelprang and Lemaux, 2020; Rahman et al., 2023). However, the use genetic engineering to improve the economic and ecological traits of *Pinus* spp. remains limited. An improved understanding of cultivar resistance and the breeding of stable and effective cultivars through genetic engineering are crucial for the management of PWD.

Systematic analysis of the molecular mechanisms of PWN pathogenicity and the selection of resistance genes are prerequisites for genetic engineering and lay the foundation for effective proactive control methods and comprehensive prevention strategies (Bláhováet al., 2020; Mackelprang and Lemaux, 2020). Previous research and empirical evidence have revealed that PWN pathogenicity begins with its proliferation in the xylem of pine trees, which causes mechanical damage (Ichihara et al., 2000; Umebayashi et al., 2011; Yazaki et al., 2018). Additionally, PWN is often accompanied by other pathogens (such as fungi), and co-infection exacerbates the damage to pine trees (Xue et al., 2019; Jia et al., 2023). Proteases and polysaccharide enzymes are secreted by PWN, and these toxins disrupt cellular structure, interfere with normal physiological functions, and lead to tree death (Zhao et al., 2007; Faria et al., 2015). Scientists have explored resistance by *P. massoniana* and revealed several key genes related to PWN resistance. Among them, *PmNBS-LRR97*, *PmGPPS1*, *PmTPS4*, and *PmTPS21* participate in defense against PWN (Xie et al., 2023; Liu et al., 2021; Liu et al., 2022). These results indicate that the related resistance genes have significant advantages in long-term pest control. However, the lack of reference genome for *P. massoniana* and a focus on single genes or single-level interactions can inhibit an overall understanding of multidimensional and multivariate relationships, reducing effectiveness in the complex forest environment over the long term.

Previously, we analyzed the spectral performance in the field of the germplasm resource library concurrent with PWN carrying capacity to identify resistant cultivars (RC) and tolerant cultivars (TC) (Liu et al., 2023a). Building on this, we constructed a full-length transcriptome library for *P. massoniana* through iso-seq, analyzed the transcriptomes and metabolomes of the RC and TC, and explored the molecular basis of *P. massoniana* resistance to PWN. An in-depth exploration of the biochemical interactions between different cultivars and PWN enabled us to clarify the strategies used by *P. massoniana* in response to PWN invasion and to determine their molecular basis. Our findings are of great significance for understanding PWD transmission and control.

## 2 Materials and Methods

### 2.1 Plant material and growth conditions

The experimental subjects were taken from the nursery at Guanzhuang State-owned Forest Farm in Sanming, Fujian, China (26.5603°N, 117.7455°E), A total of 71 clonal interforest cultivars were obtained. The study area comprised a 76 ha, 16-year-old pure *P. massoniana* plantation with an annual average temperature of 19.9 °C and precipitation of 1375.2 mm. In the early stage, three types of clones were screened based on spectral characteristics and PWN carrying amounts (PCA): tolerant cultivars (TC) (high PCA, spectral data biased towards health, low mortality), resistant cultivars (RC) (extremely low PCA, spectral data biased towards health, low mortality) and neutral cultivars (NC) (high PCA, spectral data biased towards death, high mortality) (Liu et al., 2023a). Eight clones (TC numbers 43, 45, and 58; RC numbers 52 and 59; and NC numbers 64, 41, and 51) were measured in the laboratory.

In the above *P. massoniana* cultivars, PCA and mortality were quantified 6 weeks after inoculation with PWN. Cultivars were placed in a growth chamber (16 h light, 8 h dark, 70% humidity, 28 °C in the light, and 24 °C in the dark) for 2 months before experimental treatment.

### 2.2 Pine wood nematode inoculation assay

Pine wood nematode adults were cultured as previously described (Cai et al., 2021). Seedlings of *P. massoniana* were inoculated with 1 mL of a PWN solution (5000 individuals / mL), with 15 replicates (individual plants) per cultivar. Double-distilled water served as a negative control.

### 2.3 Quantification of relative PWN levels

Real-time quantitative polymerase chain reaction (RT-qPCR) was used to quantify PWN in *P. massoniana* cultivars. Fresh branches (with leaves) collected from the entire young seedling from the laboratory, were used for PWN quantification. Total genomic DNA was extracted from each plant sample using a MoBio PowerPlant® Pro DNA Isolation Kit (cat no.12855-50, MoBio, USA), according to the manufacturer’s protocol. DNA quantity and quality were measured with a NanoDrop 2000 photometer (Thermo Fisher Scientific, USA), and DNA integrity was determined by 1% agarose gel electrophoresis. The extracted DNA was stored at -80 °C until further use. Quantitative PCR was conducted using a PWN-specific BxCW primer (F: 5 ’-TTGCATTCTACGGCCAGTCC-3’; R: 5′- ACTGACTTTCGATGGCTCCG-3’) and a host-specific PmEF2 primer (F: 5’- CTGCGATGTCCCTCATGTTA-3’; R: 5′- AACAAGGTCTTTCCCCTCGT-3’) (one transcript from *P. massoniana* was used as the internal control) (Xie et al., 2023), with Hieff™ qPCR SYBR Green Master Mix (Low Rox Plus, cat no. 11202ES08, Yeasen, Shanghai, China) on a QuantStudio 6 Flex PCR system (ABI). The qPCR signals were normalized to those of the reference gene PST in *Pinus* by applying the 2^-ΔΔCT^ method (Fang et al., 2023). Three biological replicates and three technical replicates were used. One-way analysis of variance (ANOVA; Tukey test) was used to determine differences among groups.

### 2.4 Generation of full-length reference transcripts

The sample preparation, as described in section 2.1, was repeated using three techniques with six weeks after infection by PWN. Total RNA was extracted from the cultivars using TRIzol (Invitrogen, CA, USA), eukaryotic mRNA was enriched with magnetic beads with Oligo(dT), and mRNA was reverse transcribed into cDNA using a SMARTer™ PCR cDNA Synthesis Kit. BluePippin was used to screen full-length cDNA fragments (Sage Science, Beverly, MA, USA) to generate cDNA libraries of different lengths (1–2 k, 2–3 k, and 3–6 k). The SMRT bell library was obtained by terminal repair of full-length cDNA and a SMRT dumbbell connector. The libraries were then sequenced via a PacBio RS II platform (Menlo Park, CA, USA). Trimmomatic (v0.33) was used for quality control and filtering of raw reads. Low quality reads and those with short read lengths were removed. Nonchimeric full-length sequences were identified from reads of inserts based on the poly (A) tail signals, 5′ adapter sequences, and 3′ adapter sequences. An isoform-level clustering algorithm was used to iteratively cluster the full-length transcript sequence (Chin et al., 2013). In combination with non-full-length sequences, consistent sequences were corrected using a corrective (polish) procedure. Finally, sequence redundancy was removed using CD-HIT (http://weizhongli-lab.org/cd-hit/) (Li and Godzik, 2006).

### 2.5 RNA-seq and analysis

The sample preparation, as described in section 2.4. The Illumina cDNA library was prepared using a NEBNext Ultra II RNA Library Prep Kit (NEB, Ipswich, MA, USA) according to the standard experimental protocol. An Illumina Novaseq 6000 with a PE 150 sequencing strategy was used for computer sequencing. Read segments sequenced by Illumina were mapped to SMRT cDNA libraries by Bowtie2 (Langdon, 2015) and quantified by RSEM (Li et al., 2015) using fragments per million mapped read segments (FPKM) per kilobase transcript. The Non-Redundant Protein Sequence Database (NR) (ftp://ftp.ncbi.nlm.nih.gov/blast/db/), Gene Ontology (GO) database (http://geneontology.org/), Swiss-Prot Protein Sequence Database (SwissProt) (http://www.ebi.ac.uk/uniprot), clusters of euKaryotic Orthologous Groups (KOG) tool (ftp://ftp.ncbi.nlm.nih.gov/pub/COG/), and the Kyoto Encyclopedia of Genes and Genomes (KEGG) database (http://www.genome.jp/kegg/) were used for analyses. DESeq (1.10.1) (Love et al., 2014) was used for differential expression analysis, and R software was used for correlation analysis and drawing heat maps.

### 2.6 Metabolomic analysis

Metabolites were extracted from each sample as previously described (De Vos et al., 2007). Ultra high performance liquid chromatography (UHPLC) separation was performed using an ACQUITY UPLC BEH amide column (1.7 μm, 2.1 mm × 100 mm, 186004801, Waters, Milford, USA) equipped by Oebiotech Company (Shanghai, China). The flow rate of the mobile phase was set at 0.5 mL / min and consisted of a 0.1% formic acid solution (A) (A117-50, Thermo Fisher Scientific, Waltham, MA, USA) and a 0.1% formic acid in acetonitrile (B) (A998-4, Thermo Fisher Scientific, Waltham, MA, USA). The column temperature was 25 °C, the auto-sampler temperature was 4 °C, and the sample volume was 1 μL (Chen et al., 2007; Theodoridis et al., 2008). Data were pruned from different samples to distinguish the metabolites in different resistant cultivars of *P. massoniana*. Next, commercial databases, including the KEGG database and MetaboAnalyst (https://www.kegg.jp/), were used with the search term “metabolite pathway” (https://www.genome.jp/kegg/pathway.html).

## 3 Results

### 3.1 Nomination of potential PWN-determining genes in *P. massoniana* using a full-length transcriptome

To uncover the pathogenic mechanisms of the PWN in its host pine species, we sequenced the transcriptome of *P. massoniana* and screened for potential PWN-determining genes (β) by comparing protein information from *Pinus densiflora* and *Pinus tabuliformis* and predicting the function of β. The sequencing process involved extracting unigenes from raw data through steps that included filtering, quality control, classification, clustering, correction (polishing), and redundancy removal, leading to the prediction of 55,724 coding DNA sequences (**Figure 1A**). The integrity of the transcript data across three generations was evaluated using BUSCO analysis, which revealed that 93.5% of the isoforms were complete (**Supplementary Figure S1**). Furthermore, 52,731 (96.63%) genes were annotated in at least one database (KOG, NR, KEGG, or Swissprot, among others) (**Figure 1B**). To identify potential PWN-determining genes in *P. massoniana*, 7,497 genes (α) shared between *P. massoniana* and *P. densiflora*, both of which host PWN, were screened for amino acid sequence similarity greater than 90%. Subsequently, α was compared with a non-PWN host, *P. tabuliformis*, for amino acid sequence similarity less than 90%, which resulted in the identification of 1,938 potential PWN-determining genes (β) in *P. massoniana* (**Figure 1C**). KEGG analysis of β in *P. massoniana* revealed their predominant enrichment in MAPK signaling and plant-pathogen interaction pathways. (**Figure 1D**).

**FIGURE 1.**
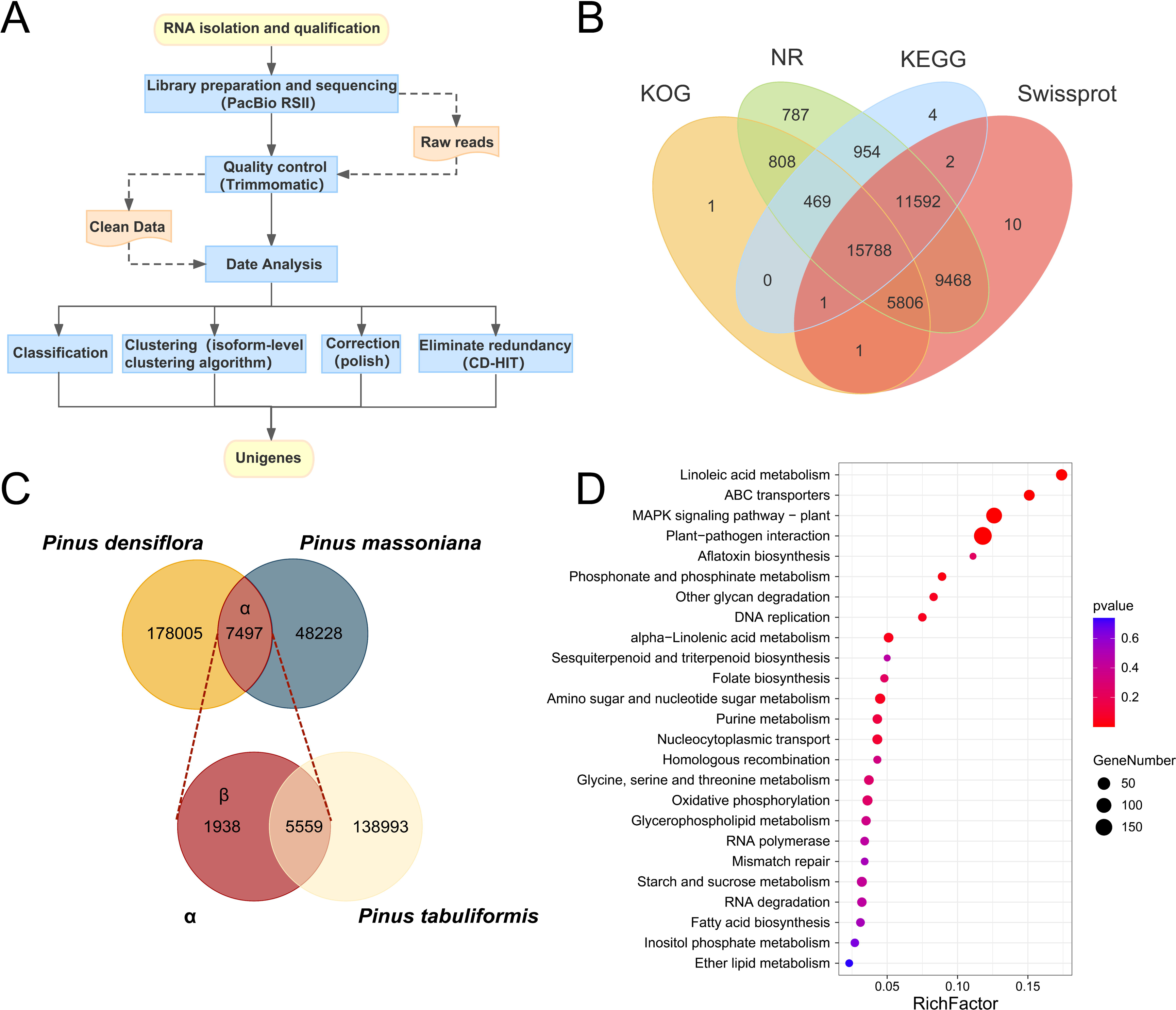
Nomination of potential PWN carrying determination genes in *Pinus massoniana* using full-length transcriptome. (**A**) Workflow of full-length transcriptome sequencing and analysis. (**B**) Venn diagrams of unigenes annotated in different databases. (**C**) To screen the potential PWN-carrying determination genes in *Pinus massoniana* (β), the common genes (α, >90% similarity of amino acid sequences) between two PWN host pine (*Pinus massoniana* and *Pinus densiflora*) were compared with proteins of non-PWN host pine (*Pinus tabuliformis*) to develop the library of non-common (β, <90% similarity of amino acid sequences) genes. (**D**) A bubble plot diagram presents the KEGG annotation of genes in β. The top 25 KEGG terms are shown.

### 3.2 Transcriptome analysis revels PWN carrier determinant genes in *P. massoniana*

Previously, the TC, NC, and RC were identified within the germplasm bank of *P. massoniana*. In this research, all three cultivars were uniformly inoculated with PWN, and the PCA were quantified six weeks post-inoculation. We found that the PCA in the TC was significantly higher than that in the NC, which in turn exhibited a higher PCA than the RC (*P* ≤ 0.01). Specifically, the PCA in the TC and RC were 1.68 and 0.34 times that of NC, respectively (**Figure 2A**). Concurrently, we identified genes that were strongly associated with the PCA (*P* ≤ 0.01, |r| ≥ 0.8), resulting in a total of 7,710 genes (**Supplementary Table S1**). These genes were then compared with β to determine the PWN carrier determinant genes (γ) in *P. massoniana*, which amounted to 318 genes (**Figure 2B**). In the TC vs NC and the RC vs NC, 128 and 63 genes were upregulated, respectively, and 116 and 200 genes were downregulated, respectively (**Figure 2C**), indicating that spatial variation in the expression of related genes occurred during PWN invasion of the TC and RC. Furthermore, KEGG functional enrichment analysis was performed to predict the function and pathway of γ and revealed significant enrichment in MAPK signaling and plant-pathogen interaction pathways (**Figure 2D**).

**FIGURE 2.**
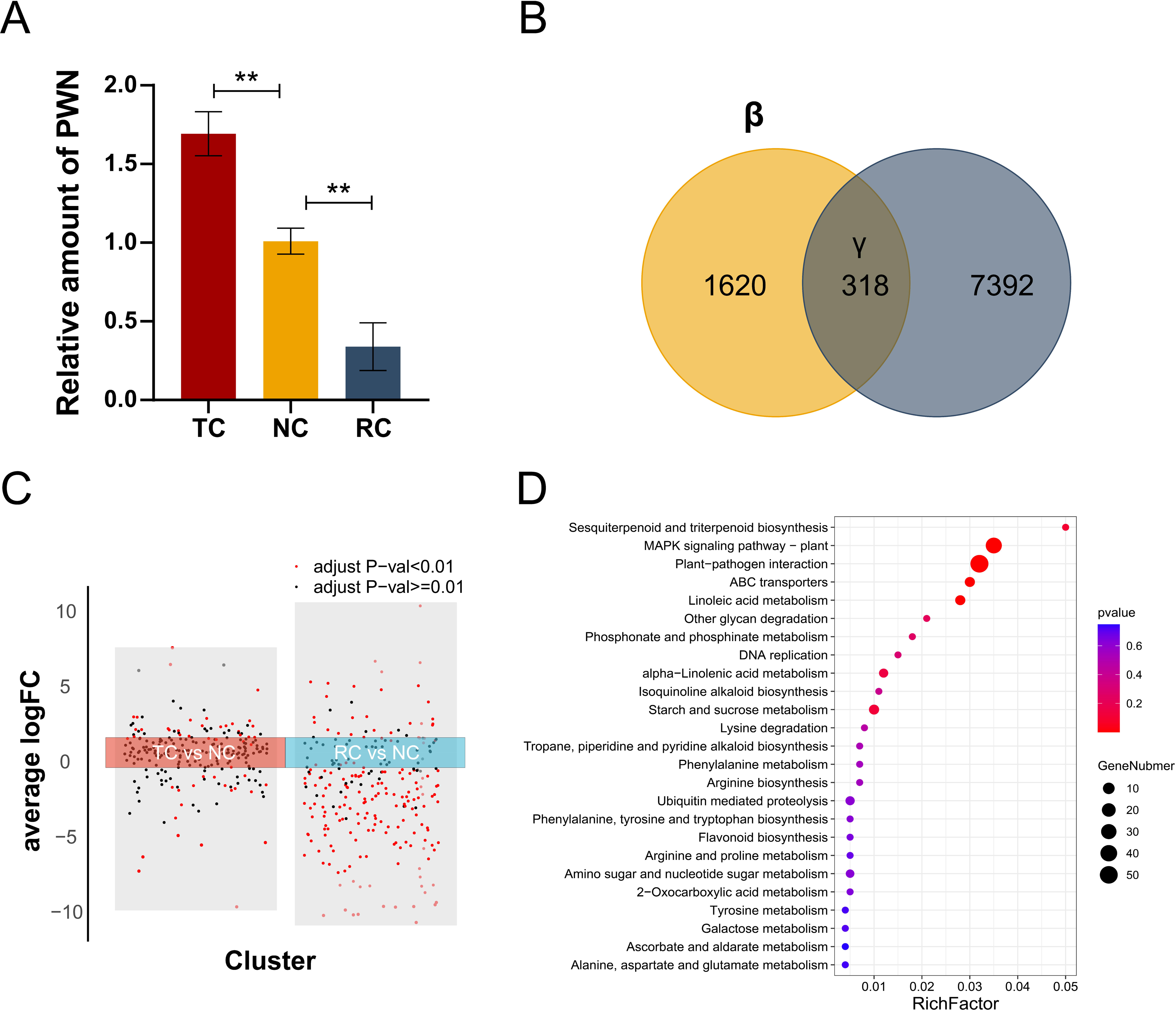
Transcriptome analysis revels the PWN carrier determine genes in *P. massoniana*. (**A**) Relative PWN carrying amount of different *P. massoniana* cultivars (TC, tolerant cultivar; NC, neutral cultivar; RC, resistant cultivar) were quantified six weeks post inoculation. Each box represents a mean value, with bars representing one standard deviation (SD). ** indicate statistical differences (*P ≤ 0.01*) between samples using one-way ANOVAs (Tukey’s test). (**B**) Venn diagrams shown the number of potential PWN-carrying determination genes in *P. massoniana* (β), differentially expressed genes (DEGs) significantly correlated with PWN amount, and PWN-carrying determination genes in *P. massoniana* (γ). (**C**) Differential gene expression analysis revealed 318 genes (γ) up-regulated and down-regulated in all two clusters. Adjusted *P ≤ 0.01* are shown in red, and adjusted *P ≥ 0.01* are shown in black. (**D**) A bubble plot diagram presents the KEGG annotation of genes in γ. The top 25 KEGG terms are shown.

### 3.3 Molecular strategies of *P. massoniana* in response to PWN invasion

Interforest spectral data previously revealed three distinct types of *P. massoniana* with varying health (Liu et al., 2023a). Consequently, we evaluated the mortality rates of these host plants six weeks after inoculation with PWN. The mortality rate of the *P. massoniana* NC was significantly higher than that of the TC or the RC (*P* ≤ 0.01); mortality was 7.4 times or 6.6 times higher in the NC, respectively (**Figure 3A**). For the PCA of each cultivar (**Figure 2A**), we found that the TC exhibited a high PCA yet maintained a low mortality rate, whereas the RC demonstrated both low mortality and PCA. We speculated that these were two different PWN resistant cultivars. After full-length transcriptome sequencing, we performed second-generation transcriptome sequencing for the three cultivars six weeks after PWN inoculation to assess gene expression in each cultivar and ascertain the molecular mechanisms underpinning resistance to PWN by the TC and RC. We identified 274 common related genes (CRGs), 243 tolerance-related genes (TRGs), and 227 resistance-related genes (RRGs) across the β in theTC vs NC, and RC vs NC comparison groups (**Figure 3B**). Concurrently, we evaluated the relative expression of the three gene categories in each cultivar (**Figure 3C**). Among them, 49% of TRGs (118 of 243) exhibited higher relative expression in the TC compared to the other two cultivars. The relative expression of RRGs in the RC was higher in 59% (161 of 274) and 71% (161 of 227) than in the other two cultivars. Additionally, KEGG enrichment analysis of CRGs, TRGs, and RRGs revealed enrichment not only in linoleic acid metabolism, MAPK signaling, and plant-pathogen interaction pathways but also in other pathways. Specifically, TRGs were enriched in the spliceosome pathway (**Figure 3D**), CRGs were enriched in ABC transporter and starch and sucrose metabolism pathways (**Figure 3E**), and RRGs were enriched in ABC transporter pathways (**Figure 3F**).

**FIGURE 3.**
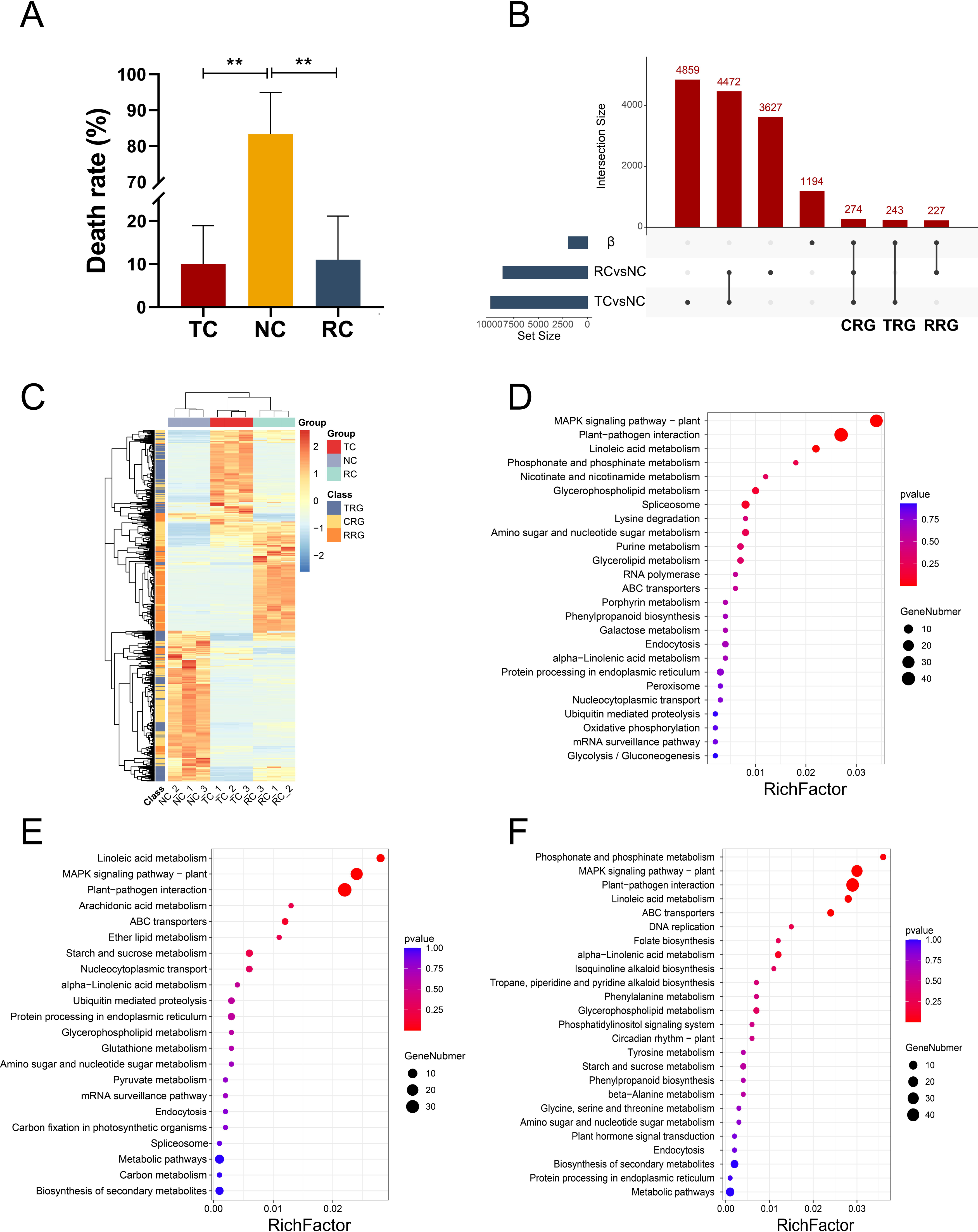
Different molecular strategies of *Pinus massoniana* in response to PWN invasion. (**A**) Mortality of different *P. massoniana* cultivars were documented six weeks post inoculating with PWN. Each box represents a mean value, with bars representing one standard deviation (SD). ** indicate statistical differences (*P* ≤ 0.01) between samples using one-way ANOVAs (Tukey’s test). (**B**) Upset maps shown the number of potential PWN-carrying determination genes in *Pinus massoniana* (β), DEGs of tolerance and neutral cultivars (TC vs NC), DEGs of resistance and neutral cultivars (RC vs NC), Common Related Genes (CRGs), Tolerance Related Genes (TRGs) and Resistant Related Genes (RRGs) (**C**) Heat maps of relative expression level of TRGs, CRGs and RRGs in different *P. massoniana* cultivars, data were homogenized by Z-score. Bubble plot diagrams present the KEGG annotation of TRGs (**D**), CRGs (**E**), and RRGs (**F**), respectively. The top 25 KEGG terms are shown.

### 3.4 *P. massoniana* metabolic responses to PWN invasion

Identifying downstream metabolites is essential to understanding *P. massoniana* resistance to PWN. Non-targeted metabolomic sequencing was performed on three cultivars six weeks after PWN inoculation. Metabolites that were highly correlated with PCA were selected in different cultivars (*P* ≤ 0.01, |r| ≥ 0.8), resulting in the identification of 465 metabolites (δ) (**Figure 4A**). Among these, 145 metabolites were positively correlated with PWN (*P* ≤ 0.01, 1 ≥ r ≥ 0.8), while 320 were negatively correlated with PWN (*P* ≤ 0.01, 1 ≤ r ≤ 0.8) (**Figure 4A**). KEGG enrichment analysis indicated that δ were significantly enriched in the arachidonic acid, arginine, and proline metabolism pathways (**Figure 4B**). Moreover, we analyzed the differential accumulation of metabolites in the TC vs NC and RC vs NC and identified a total of 302 common related metabolites (CRMs), 531 tolerance-related metabolites (TRMs), and 222 resistance-related metabolites (RRMs) (**Figure 4C**). The expression of each type of metabolite in each cultivar was calculated (**Supplementary Figure S2**). KEGG enrichment analysis of CRMs, TRMs, and RRMs showed that TRMs were primarily enriched in purine metabolism, flavone and flavonol biosynthesis, and ABC transporter pathways (**Figure 4D**). CRMs were enriched in phenylpropanoid biosynthesis, purine metabolism, and alkaloid biosynthesis pathways (**Figure 4E**). RRMs were enriched in ABC transporter and metabolic pathways (**Figure 4F**).

**FIGURE 4.**
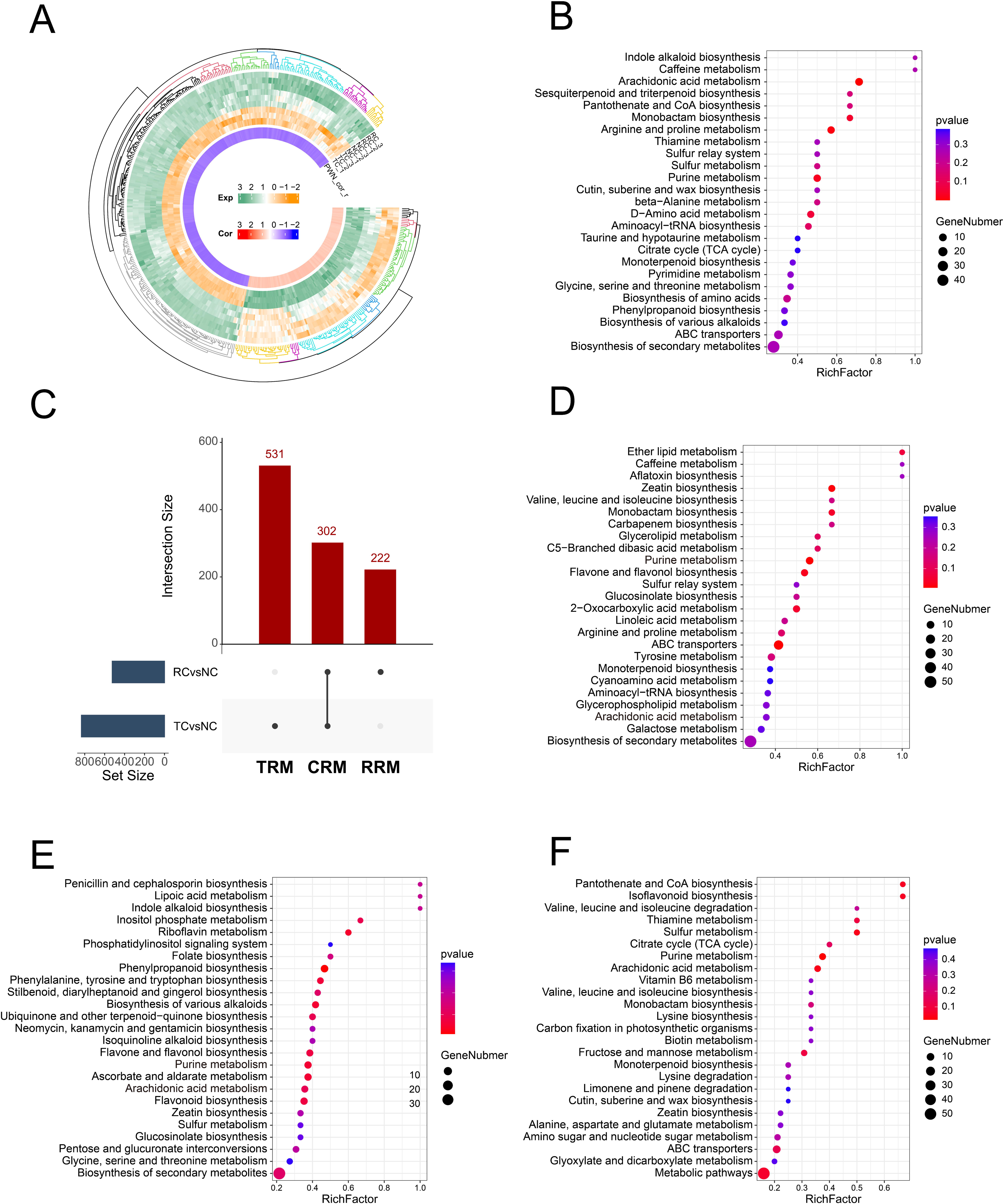
Different metabolic strategies of *Pinus massoniana* in response to PWN invasion. (**A**) The circular heat map shows the relative expression (inner circle) and correlation coefficients (outside circle) of metabolites highly correlated with PWN (δ) in different *P. massoniana* cultivars, data were homogenized by Z-score. (**B**) A bubble plot diagram presents the KEGG annotation of metabolites in δ. The top 25 KEGG terms are shown. (**C**) Upset maps shown the number of different accumulation metabolites (DAMs) in *P. massoniana* of tolerance and neutral cultivars (TC vs NC), DAMs of resistance and neutral cultivars (RC vs NC), Common Related Metabolites (CRMs), Tolerance Related Metabolites (TRMs) and Resistant Related Metabolites (RRMs). Bubble plot diagrams present the KEGG annotation of TRMs (**D**), CRMs (**E**), and RRMs (**F**), respectively. The top 25 KEGG terms are shown.

### 3.5 The genetic landscape of *P. massoniana* following PWN invasion

Integrating transcriptomic and metabolomic analyses can shed light on plant metabolic pathways. Differentially expressed genes (DEGs) and differentially accumulated metabolites (DAMs) may reveal the mechanisms behind plant resistance to pathogens. Our analysis of the transcriptomes and metabolomes of *P. massoniana* cultivars revealed genes and metabolites within the same metabolic pathways (**Supplementary Figure S3**). Based on the relative contents of pertinent genes and metabolites (**Figure 5A**), we identified three strategies by which *P. massoniana* responds to PWN invasion (**Figure 5B**). Initially, the PWN triggers a common response in all cultivars, which comprises the inhibition of endogenous hormone synthesis (IAA, ABA, aspartic acid, and cysteine), the downregulation of ABCB1 and ASP5 expression, and the upregulation of ABCC1, ABCC2, and germacrene D synthase expression. Alterations in metabolites, including L-arginine, glutamic acid, myo-inositol, (-)-riboflavin, trehalose, and aspartic acid, ultimately restrict the reproductive and pathogenic processes of the PWN. In contrast to the NC, the RC relies on specialized mechanisms to curtail PWN reproduction. Upon PWN invasion, the RC initially promotes the accumulation of linoleic acid and induces the synthesis of JA through AOC, AOC3, and LOX2S (lipoxygenase) proteins, thereby regulating downstream genes (EGL3, MYC2) and metabolites. Pine wood nematode promotes the accumulation of ROS and the synthesis of chitinase but also limits the amount of PWN carried by *P. massoniana*, which protects the tree from PWD. The TC also employs special mechanisms to limit PWN pathogenicity. Initially, PWN invasion triggers the accumulation of DPPC (phosphatidylcholine), which, under the action of LPIN (phosphatidate phosphatase), stimulates the synthesis of linolenic acid. This regulates the production of JA through LOX1<u>_</u>5 (linoleate 9S-lipoxygenase) and LOX2S proteins. Jasmonic acid then promotes the synthesis of downstream flavonoid metabolites (neohesperidin dihydrochalcone, flavonol base, flavone base, DL−phenylalanine, and neobavaisoflavone), which limits the pathogenicity of the PWN and protects *P. massoniana* from PWD.

**FIGURE 5.**
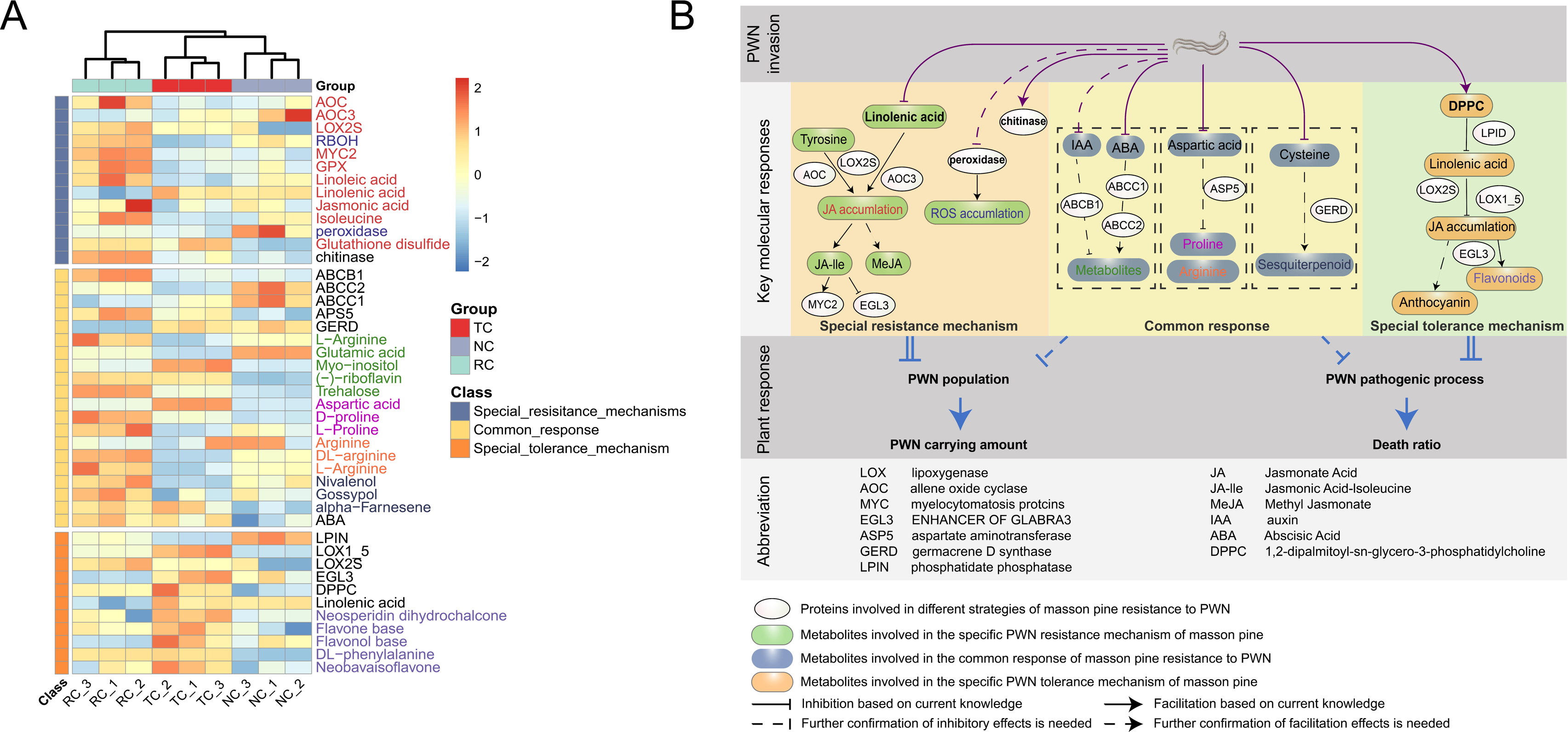
Genetic landscape in *Pinus massoniana* regard to PWN invasion. (**A**) Heat map represent the relative expression level of key different expressed genes (DEGs) and different accumulation metabolites (DAMs) involved in different strategies of *P. massoniana* against PWN invasion, data were homogenized by Z-score. (**B**) Illumination of critical genetic responses in *P. massoniana* against PWN invasion regard to different ecological cultivar

## 4 Discussion

As the primary host for PWN in southern China, it is important to understand how *P. massoniana* responds to PWN invasion. Indoor tests confirmed that PCA and mortality were low in the RC that was screened from the forest, while the TC had a high PCA but low mortality. On this basis, we explored broad-spectrum resistance to PWN and the molecular basis for resistance and tolerance in *P. massoniana* via transcriptomic and metabolomic analyses (**Figure 5B**).

Initially, we conducted RNA-seq on three cultivars using the reported full-length transcriptome of *P. massoniana* as a reference, but the mapping rate for the second-generation transcriptome was low (Liu et al., 2020). Therefore, we performed full-length transcriptome sequencing on 71 varieties in the seed orchard and obtained 55,724 high-quality transcripts with an average length of 3,343.56 bp, and 93.5% of the transcripts were annotated in the database. Using this library as a reference genome, the average splicing ratio of the second-generation transcriptome sequencing results reached 94.73%. Previously, 81,837 high-quality transcripts with an average length of 846.54 bp were reported, and the average splicing ratio for second-generation transcriptome sequencing was 85.02% (Liu et al., 2020). The full-length transcriptome sequencing results in this study were superior to previous data in several respects, with the exception of the number of transcripts. However, considering the average length of transcripts, the volume of data in this study is larger than that for reference transcripts in previous studies. Additionally, not all pine trees are hosts for the PWN; therefore, we screened for potential PWN interaction-related genes in *Pinus* by using comparative genomics. According to genome data for *P. densiflora* (another PWN host) and *P. tabuliformis* (a non-host), 1,938 common potential PWN-determining genes (β) differed significantly in host trees relative to non-host trees. These were defined as potential PWN-host pine tree interaction proteins and were enriched in the MAPK signaling and plant-pathogen interaction pathways. The MAPK signaling pathway is involved in two major barriers, pathogen-associated molecular pattern (PAMP)-triggered immunity and effector-triggered immunity, which help plants resist microbial invasion. By transmitting and amplifying external signals, specific defense genes are activated and promote plant resistance (Manna et al., 2023; Peng et al., 2023). The plant-pathogen interaction pathway involves the recognition of specific molecular patterns that trigger immune responses and prevent microbial invasion and spread (Kunkel and Johnson, 2021; Rubio-Somoza and Blázquez, 2023). These findings suggest that the formation of PWN hosts is related to associated microbes.

In field experiments, over 87% of *P. massoniana* cultivars produced a common response but were unable to restrict the proliferation or pathogenesis of the PWN, which corroborates observations of severe, widespread PWD in the field. The common response of *P. massoniana* included the ABC transporter, arginine, and proline pathways. First, ABC transporters were involved in the transport of secondary metabolites and plant hormones, detoxification, and signal transduction, which indirectly aided resistance to PWN (Wang et al., 2022; Li et al., 2024; Zhou et al., 2024). Second, arginine and proline indirectly affected the reproduction and pathogenicity of the PWN by promoting plant growth and development, enhancing stress resistance, acting as antioxidant defense molecules, participating in defense responses, and regulating metabolic pathways (Zhang et al., 2022; Theisen et al., 2024; Wang et al., 2024b). The specific mechanisms by which ABC transporters, arginine, and proline function in the resistance of *P. massoniana* to PWN require further study. In the common response of *P. massoniana*, monoterpenes and sesquiterpenes had multiple roles as chemical defense substances. First, they had a direct toxic effect on the PWN, inhibiting its growth and reproduction by disrupting its nervous system or metabolic pathways (Hwang et al., 2021, 2022; Sánchez-Martínez et al., 2021). Second, these substances can act as signaling molecules that induce plants to produce defense enzymes and secondary metabolites that enhance resistance (Castorina et al., 2020; Paul et al., 2020; Xu et al., 2021). The antioxidant effects of monoterpenes and sesquiterpenes also alleviate oxidative stress caused by PWN infection (Farhadi et al., 2020; Son et al., 2023). However, the common response is insufficient in limiting the PWN, possibly because it may detoxify secondary metabolites (including terpenoid compounds) (Jia et al., 2023). In future studies, adding monoterpenes and sesquiterpenes to the diet of the PWN and observing mortality may indicate whether the PWN has a detoxification system.

In this study, chitinase, the JA signaling pathway, and related DEGs and DAMs involved in maintaining ROS homeostasis were enriched in the RC and served as the main effector signaling pathways for defending against the PWN (**Figure 5A**). Stress may disrupt ROS homeostasis, leading to ROS accumulation, which in turn promotes the accumulation of JA (Wang et al., 2019; Furuta et al., 2024; Shi et al., 2024) and upregulates the expression of chitinase, ultimately affecting the PCA (Shen et al., 2022; Qiu et al., 2023). The role of chitinase in protecting plants from stress has been well established (Ju et al., 2016; Berini et al., 2018; Jha and Modi, 2018; Lu et al., 2020); chitinase has been shown to degrade chitin in nematode eggshells, which limits nematode reproduction (Chan et al., 2010; Lee and Kim, 2015; Chen et al., 2021). This strategy has been widely applied in the control of agricultural nematodes. Therefore, we speculate that in the RC, chitinase expression is the main mode of action of resistance (**Figure 5B**). In future studies, verifying the direct effect of chitinase on the PWN will allow us to ascertain its importance in defense and its potential as a target for new eco-friendly nematicides (Zhao et al., 2023).

Flavonoids play a key role in tree defense against the PWN through signal transduction and metabolic regulation (Isshiki et al., 2014; Vazquez-Vilar et al., 2023; Wang et al., 2024a). Following PWN invasion, the host recognizes PWN secretions and related microbial products, activates defense signaling pathways, and induces the expression of flavonoids (Park et al., 2020). These substances have antioxidant and antimicrobial functions that limit PWN pathogenicity and microbial infection (Raorane et al., 2019; Liu et al., 2023b). Pine wilt nematode pathogenicity involves numerous symbiotic or associated microbes (Jia et al., 2023). Flavonoids affect the environment for nematodes by inhibiting the growth of certain pathogenic microbes or regulating the structure of beneficial microbial communities (Borah et al., 2018). Therefore, the role of flavonoids in PWN invasion is not limited to direct defense but also involves regulating complex interactions between the host plant and microbes. We found that the tolerance strategy promoted synthesis of the JA substrate linolenic acid, which increased the accumulation of flavonoids (**Figure 5B**). The TC had the highest flavonoid content (**Figure 5A**), suggesting that it regulates PWN-associated microbes by accumulating flavonoids and limiting PWD, which may be the reason for low mortality in the TC (**Figure 3A**).

The molecular strategies of all three varieties of *P. massoniana* cultivars involved the JA signaling pathway. However, due to the unique genetic backgrounds of *P. massoniana* cultivars, there were significant differences in the composition and regulation of the JA signaling pathway. The RC may possess more sensitive JA receptors or more effective signal transduction components, thereby exhibiting higher resistance to PWN invasion (**Figure 1A**). It is worth noting that the JA signaling pathway does not exist in isolation but interacts with other plant hormones (such as auxin and abscisic acid) and environmental signals (such as light and temperature) to form a complex regulatory network (Qiu et al., 2022; Hu et al., 2023b; Sybilska and Daszkowska-Golec, 2023). Different pines may exhibit different regulatory interactions that affect their defense mechanisms. Over the long term, different pines may have developed specific defense mechanisms under different environmental conditions (Kalachova et al., 2023). Epigenetic factors, such as DNA methylation and histone modification, may also affect the activation and expression of the JA signaling pathway, thereby leading to differences in tree defense (Campos-Rivero et al., 2017; Du et al., 2024). Subsequent studies can investigate the impact of different exogenous signals (such as plant hormones or environmental changes) on *P. massoniana* resistance in the field. By exploring the interactions between different signaling pathways, we can better understand *P. massoniana* defense in the natural environment and provide a scientific basis for the management of PWD.

In summary, transcriptome and metabolome analyses provided new insights into the genetic diversity and genetic basis of pine tree resistance to PWN invasion. Our results clarified the mechanisms by which the RC restricted PWN reproduction through chitinase synthesis and the TC protected plant health via flavonoid accumulation. Future research on molecular interactions between different resistant cultivars and the PWN, the effects of chitinase and flavonoids, and the mechanisms of the response process will further expand our understanding of PWD and its prevention and control.

## 6 Author Contributions

JS designed and managed the research. JW and SC wrote the manuscript. LZ and JT performed all analyses and drawings. DH, XZ and TF conducted the field test and sampling. SS, JJ and CL designed the chemical treatment related experiments.

## 7 Acknowledgments

We are very grateful to the Fujian Forestry Bureau for its help providing detailed information of sample areas in this study.

